# Structural brain architectures match intrinsic functional networks and vary across domains: A study from 15000+ individuals

**DOI:** 10.1101/2019.12.17.879502

**Authors:** Na Luo, Jing Sui, Anees Abrol, Jessica A. Turner, Eswar Damaraju, Zening Fu, Lingzhong Fan, Jiayu Chen, Dongdong Lin, Chuanjun Zhuo, Yong Xu, David C. Glahn, Amanda L. Rodrigue, Marie T. Banich, Godfrey D. Pearlson, Vince D. Calhoun

**Author notes:** Correspondence goes to: Vince D. Calhoun, Professor, Director, TReNDS Center, Georgia State University, 55 Park Place NE, Atlanta, GA 30303, Jing Sui, Professor, Institute of Automation, Chinese Academy of Sciences, Beijing 100190, China.

## Abstract

Brain structural networks have been shown to consistently organize in functionally meaningful architectures covering the entire brain. However, to what extent brain structural architectures match the intrinsic functional networks in different functional domains remains under explored. In this study, based on independent component analysis, we revealed 45 pairs of structural-functional (S-F) component maps, distributing across 9 functional domains, in both a discovery cohort (n=6005) and a replication cohort (UK Biobank, n=9214), providing a well-match multimodal spatial map template for public use. Further network module analysis suggested that unimodal cortical areas (e.g. somatomotor and visual networks) indicate higher S-F coherence, while heteromodal association cortices, especially the frontoparietal network (FPN), exhibit more S-F divergence. Collectively, these results suggest that the expanding and maturing brain association cortex demonstrates a higher degree of changes compared to unimodal cortex, which may lead to higher inter-individual variability and lower S-F coherence.

## Introduction

Human brain is a complex network of neurons that link physical neural structure to multiple human functions (Power JD et al. 2010; Alexander-Bloch A et al. 2013). Multiple computational studies have suggested that the underlying anatomical architecture of cerebral cortex shapes resting state functional connectivity on multiple time scales (Misic B et al. 2016). More evidence has now begun to suggest that specific networks derived from gray matter architectures are resembling intrinsic functional resting-state networks (Stephen Smith ED, Adrian Groves, Thomas E. Nichols, Saad Jbabdi, Lars T. Westlye, Christian K. Tamnes, Andreas Engvig, Kristine B. Walhovd, Anders M. Fjell, Heidi Johansen-Berg and Gwenaëlle Douaud 2019). Although structural networks of the human brain have typically been constructed directly using various white matter connectivity measurements obtained from diffusion weighted imaging (Bassett DS and ET Bullmore 2009), they have also been inferred indirectly from the inter-regional covariation of gray matter measured at the group level, providing information of spatially distinct regions with common covariation among subjects (Xu L et al. 2009). In parallel, the regions spatially correlated using time courses derived from spontaneous fluctuations at “resting brain” were identified as the intrinsic functional resting-state networks. Using these two kinds of features, Segall *et al*. revealed that basal ganglia network exhibited highest structural to resting-state functional spatial correlations on 603 healthy participants (Segall JM et al. 2012). In addition to offering information about the structure-function relationship of the healthy brain, various multimodal fusion studies also revealed that impaired structure topography are correlated with functional damages in mental disease (Luo N et al. 2018; Sui J et al. 2018). However, to date, fundamental questions have not been answered on whether different structure-function correspondence can be detected in different function domains/modules, *i*.*e*., unimodal cortex and heteromodal association cortex. In addition, identification of patterns in a larger dataset (15000+ participants) with more components can provide a finer and more common degree of details, providing a stable structure-function correspondence template that may be of use to the larger neuroimaing community.

To this end, we have used a discovery dataset of 7104 functional scans (within 6005 structural scans were matched for same subjects) collected at the University of New Mexico and the University of Colorado Boulder, and a replication dataset of 9214 participants from UK Biobank. As shown in **Figure 1**, first, 100 “source-based morphometry networks” with spatially distinct regions were identified based on independent component analysis (ICA), providing information about localization of gray matter (GM) variation and their covariation among individuals (Xu L *et al*. 2009). Similar job was done for the 7104 resting-state functional magnetic resonance imaging (fMRI) scans, generating 100 intrinsic functional networks (components). These GM and fMRI components were subsequently parcellated into 9 brain network modules. Spatial coherence were measured between the effective grey matter networks and intrinsic functional connectivity networks by spatial correlation. Second, segmentation on GM of replication data were conducted, to verify the reproducibility of the identified structural-functional (S-F) paired components. Third, the replicated S-F component pairs were further compared across different domains. Interestingly, the unimodal cortical areas (e.g. somatomotor and visual networks) indicate higher S-F coherence in both discovery and replication data, while those made from heteromodal association cortices, e.g., frontoparietal network, exhibit more S-F divergence. To the best of our knowledge, this is the first study to assess differences of structure-function coherence across different function domains on the currently largest dataset.

**Figure 1.**
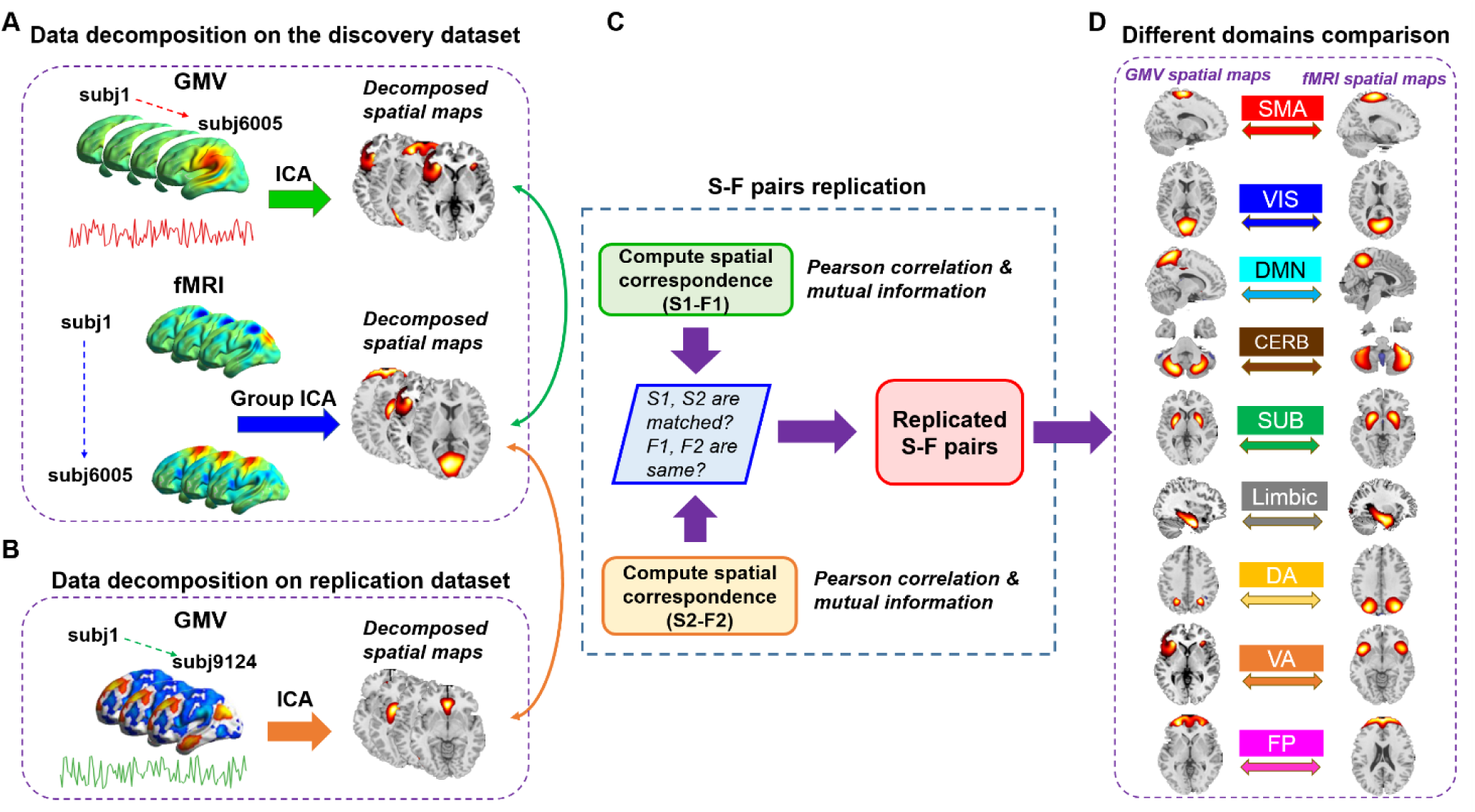
Schematic of preprocessing and analyses pipelines. (A) Structural data and functional data were preprocessed through an automated pipeline and then decomposed using spatial ICA. We then compared the correspondence between structural and functional components using Pearson correlation and mutual information. (B) A replication dataset from UK Biobank, consisting of 9214 subjects, was preprocessed to validate the identified S-F pairs. We again applied ICA to decompose the structural replication data and measured the spatial correspondence between components in the discovery dataset and components in the replication dataset. (C) If one matched S-F pair presented a high correspondence with the same functional component, then the S-F pair in the discovery dataset was regarded as replicated. (D) We further sorted the matched pairs into 9 networks and compared S-F coherence across networks.

## Materials and Methods

### Data acquisition and preprocessing

#### Discovery data

All 6101 structural scans and 7500 resting state functional scans were collected from anonymized subjects with informed consent at the University of New Mexico (UNM) and the University of Colorado Boulder (UC, Boulder). Data from the UC, Boulder site were collected using a 3T Siemens TIM Trio MRI scanner with 12 channel radio frequency coils, while data from the UNM site were acquired using the same type of 3T Siemens TIM Trio MRI scanner, and a 1.5T Avanto MRI scanner. All the data were previously collected, anonymized, and had informed consent received from subjects including both healthy and patients. As it is a de-identified convenience dataset, we do not have access to the health and identifier information. We have confirmed that the brain images do not have any obvious pathology or atrophy. The fMRI data were used in a previous study that evaluated replicability in time-varying functional connectivity patterns (Abrol A et al. 2017). The sMRI data were used in a previous study which measured age-related structural variations across the adult lifespan. The details of data acquisition and preprocessing are as bellows.

T1-weighted structural images were acquired with a five-echo MPRAGE sequence with TE = 1.64, 3.5, 5.36, 7.22, 9.08 ms, TI = 1.2 s, TR = 2.53 s, number of excitations = 1, flip angle = 7°, field of view = 256 mm, slice thickness = 1 mm, resolution = 256 × 256. The structural images were then preprocessed using voxel based morphometry (VBM) based on the SPM12 old segmentation, including: (1) spatial registration to a reference brain; (2) tissue classification into gray matter, white matter and CSF using SPM12 old segmentation; (3) bias correction of intensity non-uniformities; (4) spatial normalization to the standard Montreal Neurological Institute (MNI) space using nonlinear transformation; (5) modulated by scaling with the amount of volume changes. The modulated GM data, representing the GM volumes, were resliced to 2 mm × 2 mm × 2 mm and smoothed with a 10 mm Gaussian model (Silver M et al. 2011). The smoothed GMV was then correlated to the mean of all scans to identify outliers. Those scans with a correlation less than 0.7 were removed, thus leaving behind a total number of 6005 scans for the correspondence analysis. The demographic information of the 6005 subjects were shown in **Figure 2**.

**Figure 2.**
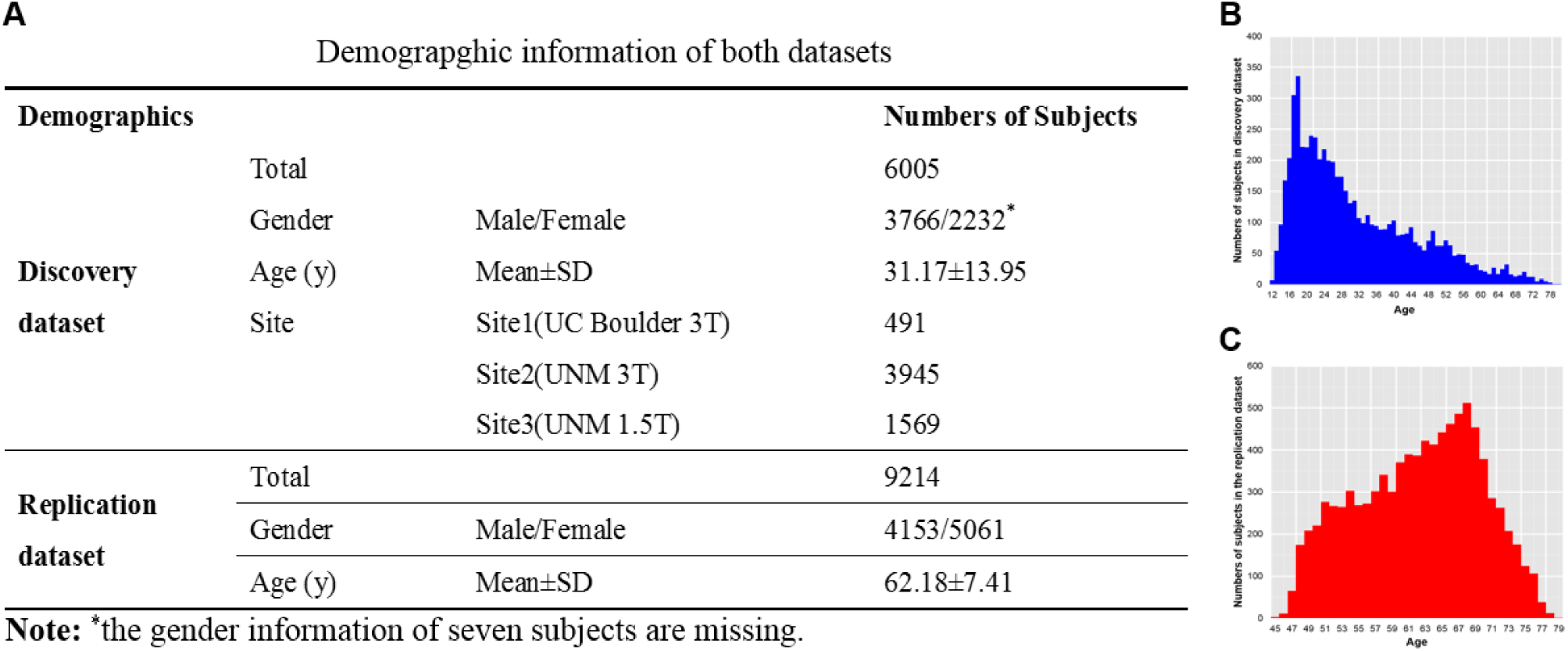
Demographic information of both datasets (A) and the distribution of age in the discovery data (B) and the replication data (C).

T2-weighted functional images were acquired using a gradient-echo EPI sequence with TE = 29 ms, TR = 2 s or 1.3 s, slice thickness = 3.5 mm, flip angle = 75°, field of view = 240 mm, slice gap = 1.05 mm, voxel size = 3.75 mm × 3.75 mm × 4.55 mm, matrix size = 64 × 64. The data preprocessing pipeline included discard of the first three images for the magnetization equilibrium, realignment using INRIalign, timing correction with the middle slice as reference, spatial normalization into the MNI space, reslicing to 3×3×3 mm and smoothing with a 10 mm Gaussian model (Silver M *et al*. 2011). More details were provided in our previous study (Abrol A *et al*. 2017). After preprocessing, 7104 functional scans were remained for the subsequent analysis, of which 6005 scans have corresponding structural images.

#### Replication data

The UK Biobank is a large-scale prospective study of over 500,000 individuals from across the United Kingdom, with a major aim being to characterize subjects before disease onset. Participants were 40-69 years of age at baseline recruitments. Here, we used the sMRI data from the February 2017 release of ∼10,000 participants (Alfaro-Almagro F et al. 2018). VBM-related processing was performed with FMRIB Software Library v10.0. A study-specific template was created using an average T1-weighted image (provided by the UK Biobank) from 5,000 subjects. To generate the template, brain extraction and tissue segmentation was performed on the average T1-weighted image. The gray matter image from the segmentation was then registered to the avg152T1_gray template available in FSL. Segmented gray matter images from each subject, available as part of the UK Biobank imaging data release, were non-linearly registered to the study specific template. Each registered grey matter image was also multiplied by the Jacobian of the warp field as a compensation (or “modulation”) for the contraction/enlargement due to the non-linear component of the transformation. The resulting GM image was then smoothed with a 6 mm Gaussian kernel. The smoothed GM was then correlated with the mean of all scans to remove scans with a correlation less than 0.7, resulting in a total number of 9214 subjects for the analysis. The demographic information of the replication data was shown in **Figure 2**.

### Group ICA on rsfMRI data

ICA decomposition on the fMRI data were conducted in our previous study (Abrol A *et al*. 2017) using Group ICA based on the GIFT toolbox (Calhoun VD et al. 2001), with a model order of 100 components **(Figure 1A)**. The spatial maps and time courses of the components were examined to select physiologically non-artifactual and previously established functional networks, as reported in (Allen EA et al. 2011; Du YH et al. 2015). Following this, 61 components were selected, which had local peak activations lying in gray matter, with time-courses dominated by low-frequency fluctuations, and exhibiting high spatial overlap with the established rsfMRI networks.

### Source-based morphometry

The segmented GM images were decomposed using spatial ICA through the GIFT toolbox (Xu L *et al*. 2009; Cota Navin Gupta JAT, Vince D. Calhoun 2017), which linearly decomposed the GM matrix into a mixing matrix that represents the relative weight of each subject for every component, and the source matrix representing the maximally spatially independent GM regions. We chose a model order of 100 components to match the numbers of components used in the fMRI analysis **(Figure 1A)**. All 100 structural components were visually inspected by three experts. We excluded structural components that had significant spatial overlaps with ventricles, white matter, large vasculature, and the brainstem, or components located at the boundaries between these regions and GM. For the purpose of spatial correlation, the GM components were resliced to 3 mm × 3 mm × 3 mm to match the dimensions of the functional components.

We then defined 9 domains/networks based on Yeo et al.’s seven-network template (Yeo BTT et al. 2011), with two extended networks including cerebellar and subcortical network. The 9 networks are: visual network (VIS), somatomotor network (SM), dorsal attention network (DA), ventral attention network (VA), subcortical network (SUB), limbic network (LIMBIC), frontoparietal network (FP), DMN and cerebellar network (CB). All the effective GM and fMRI components were further grouped into the 9 networks following a criteria based on which network/domain the peak region belongs to.

### Spatial cross-correlation between structural and functional components

To assess both linear and nonlinear spatial correspondence, we calculated spatial correlation between the selected structural and functional spatial maps using Pearson correlation and mutual information. Given two random variables x and y, their Pearson correlation can be defined in terms of their covariance cov(x, y), standard deviation of x and y as Equation (1) and their mutual information (**Figure 1C**, computed using the mutualinfo package in Matlab) is defined in terms of their probabilistic density functions p(x), p(y), and p(x, y) as Equation (2).

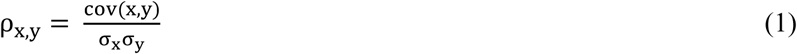

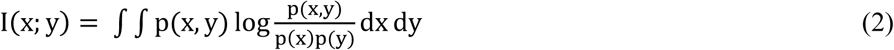

Note that before computing the correspondence, the ICA-decomposed spatial maps have been converted to Z-scores and thresholded at |Z|>2.

### Replication using the UK Biobank data

In order to validate the matched S-F component pairs, we used spatial ICA to decompose the replication data with the same model order of 100 components **(Figure 1B)**. The same inclusion criteria of component selection in the discovery dataset was applied to select good components in the replication dataset. We then computed the spatial correlation between GM components in the replication dataset and fMRI components in the discovery dataset, as well as GM components in the discovery dataset **(Figure 1C)**. If one matched structural-structural pair between discovery and replication cohorts both show high correlation with the same fMRI component, then the S-F pair in the discovery cohort was regarded as replicated.

### Comparison between different network modules

We subsequently counted the numbers of matched S-F pairs in each network module using the discovery dataset and the replicated percentage in each network module using the replication dataset. We then added up the values of PC and MI in both cohorts for each S-F pair and sorted them into a decreased order to explore which network module would present more S-F correspondence and which module indicate more S-F divergence (**Figure 1D**). Moreover, we examined the S-F correspondence of different network modules using PC or MI separately.

## Results

### Structural architectures match intrinsic functional networks

In the discovery dataset, 71 structural GM components (**Figure S1**) and 61 fMRI components (**Figure S2**) were retained for analysis after removing artifactual components through visual inspections by three professors. Out of the 71 GM versus 61 fMRI components comparisons, 44 (62%) structural components were matched with 47 (77.05%) functional components passing the predetermined Pearson correlation (PC) coefficient threshold of |r| > 0.25 and mutual information (MI) threshold of MI > 0.2 (**Figure S3**). Note that the correlation coefficient threshold corresponds to a significance level of *p*<1e-12, passing Bonferroni correlation (*p*=0.05/71/61). Empirically, we set 0.2 as threshold for mutual information and when we increased the threshold, the main results were maintained. As more than one functional component matched per structural component, and also one functional component sometimes matched with several structural components, these matched components in discovery dataset together formed 70 S-F pairs. After sorting the matched S-F pairs into 9 domains/networks, we computed the numbers of matched S-F pairs in each brain network of the discovery dataset (**Figure 3)**. The numbers of matched S-F pairs were higher in the visual network (15 pairs), default model network (13 pairs), and cerebellar network (11 pairs), but relatively lower in ventral_attention (5 pairs), dorsal_attention (5 pairs) and limbic networks (2 pairs).

**Figure 3.**
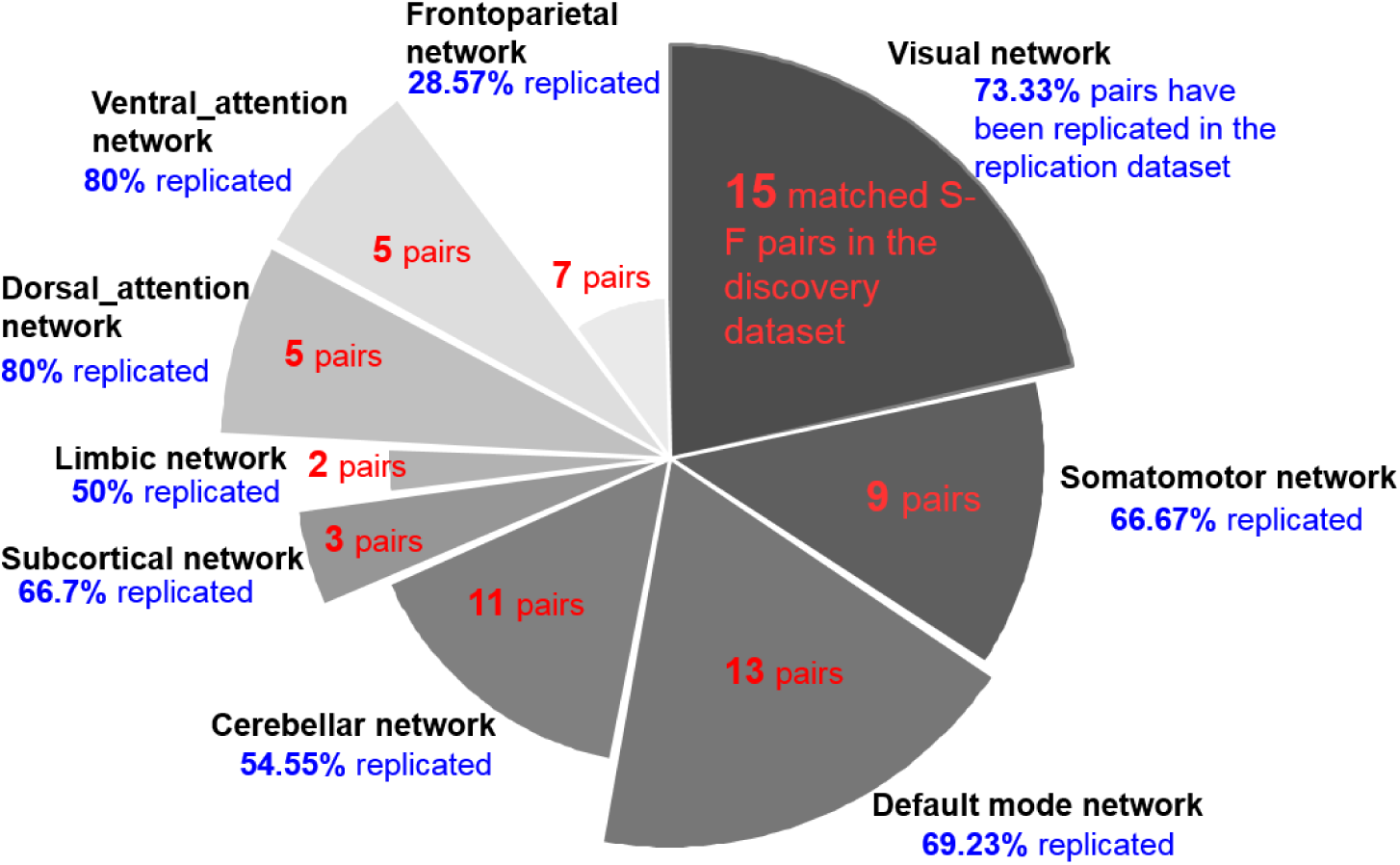
The numbers of matched pairs in the discovery data and the replicated percentages in the replication data. The S-F correspondence in visual (15 pairs), default model network (13 pairs) and cerebellar (11 pairs) networks are higher, whereas the correspondence in ventral_attention (5 pairs), dorsal_attention (5 pairs) and limbic networks (2 pairs) are relatively low. Moreover, replicated results indicate that the visual (73.33%) and DMN (69.23%) are highly replicated, while the frontoparietal network (28.57%) is not well replicated.

95 structural components in the replication data were selected as non-artifactual components after ICA decomposition. In the comparison of 95 structural components (replication dataset) and 61 functional components (discovery dataset), 66 (69.47%) structural components were matched with 49 (80.33%) functional components **(Figure S4)**. Meanwhile, 57 (60%) structural components in the replication dataset were matched with 50 (70.42%) structural components of discovery dataset **(Figure S5)**. If one matched structural-structural pair between discovery and replication cohorts both showed high correlation with the same fMRI component, then the S-F pair in the discovery cohort was regarded as replicated. In total, 45 (64.28%) out of the 70 matched S-F pairs in the discovery dataset were replicated in the UK Biobank data as shown in **Figure 4**. We set the same thresholds (|r| > 0.25 and MI > 0.2) as the discovery dataset to select the significant corresponding component pairs. The replicated percentages in each of networks are presented in **Figure 3**, which indicates that the visual (73.33%) and default mode network (69.23%) are highly replicated, while the frontoparietal network (28.57%) is not well replicated. The detail information on spatial maps and corresponding values of these 45 replicated S-F pairs are displayed in **Figure S6 to Figure S9**.

**Figure 4.**
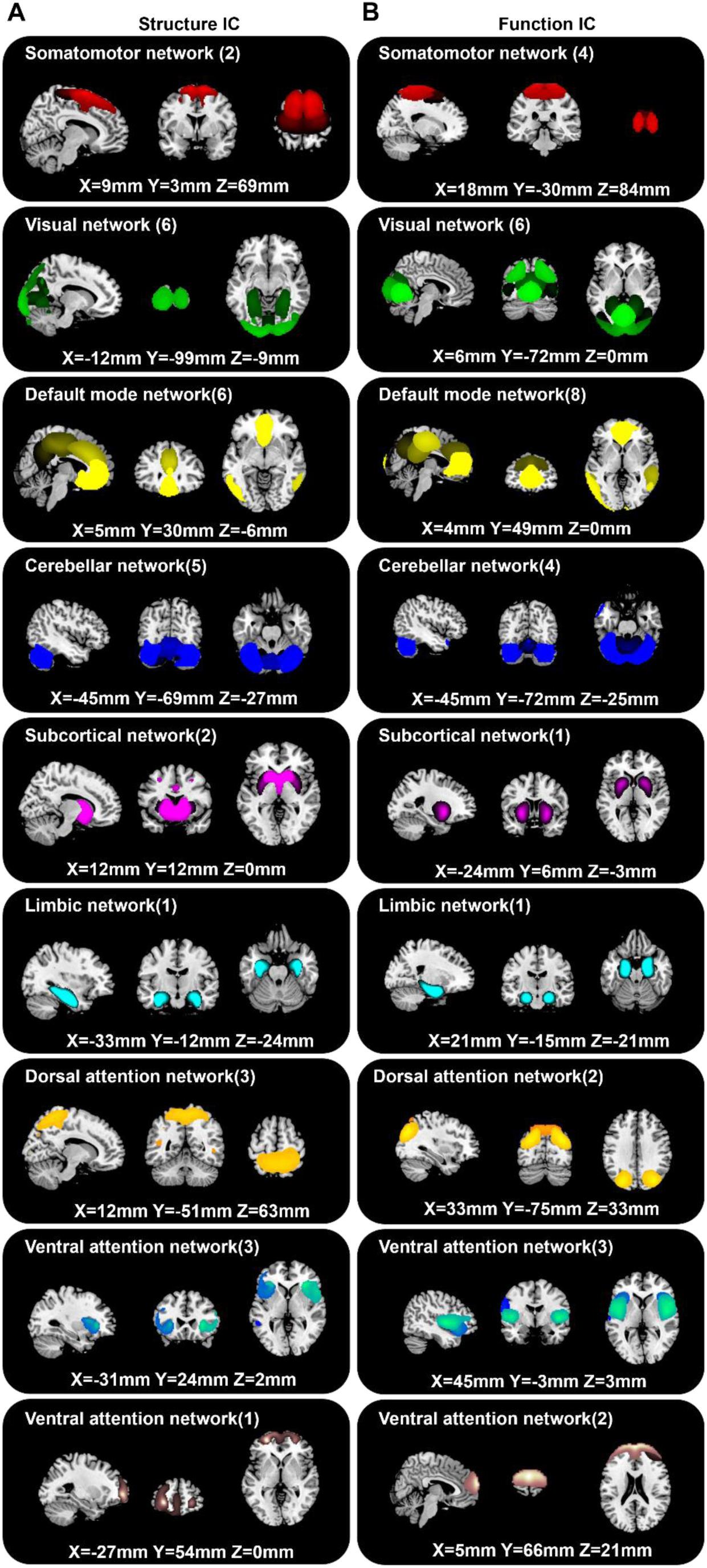
The 45 replicated matched structural (A) and functional (B) pairs in 9 networks.

### Somatomotor network

A large component, s-IC 12, spanning the supplementary motor areas and bilateral pre- and post-central gyri, are correlated with four rsfMRI components **(Figure S6A)**. The rs-IC 72, with peaks at the precentral gyri, presents the highest correspondence of PC and MI to the structural component for both discovery and replication dataset. The other three rsFMRI components are rs-IC49 and rs-IC52, centered at the paracentral lobule, and rs-IC99, which represents the bilateral postcentral gyri. A second structural component, s-IC19, which is also quite large and contains voxels spanning much of the supplementary motor area, is correlated to rs-IC36 and rs-IC13, with peaks at aspects of the supplementary motor area.

### Visual network

Notably, the visual network includes the largest numbers of S-F pairs, which is also observed in the replication dataset **(Figure 3)**. Structural component s-IC16, which largely centers at the calcarine gyrus, presents the second highest correspondence to functional component rs-IC25 in all matched S-F pairs and replicated in rep-sIC60 **(Figure S6B)**. The functional components rs-IC63 and rs-IC 96, centered at the calcarine gyrus, are also correlated with s-IC16. The other smaller structural component s-IC13 with peaks at calcarine gyrus and lingual gyrus is correlated with rs-IC25, as well as rs-IC17 and replicated in rep-sIC38. Component s-IC95 and s-IC96 comprised of a component pair, with peaks at right and left calcarine region respectively, which are correlated with rs-56, rs-IC3 and rs-IC63. Another correlated region is the lingual gyrus, where component s-IC66 is associated with rs-IC 17 and replicated in rep-sIC66. The third region with a replicated S-F correspondence in the visual network is middle occipital gyrus. Structural components s-IC 79 is correlated with functional component rs-IC76, with peaks at the bilateral middle occipital gyrus.

### Default mode network

The default mode network also shows a high S-F correspondence **(Figure S7C)**. Component s-IC31, primarily comprised of the middle cingulate gyrus, is highly correlated with rs-IC 67 and rs-IC46. Component s-IC7, which contains voxels residing in the precuneus area, is correlated with three functional components. In order of correspondence magnitude they are rs-IC34, rs-IC46 and rs-IC30, which presents aspects of precuneus and middle cingulate gyrus. Structural components, s-IC45 and s-IC52, are symmetrical components, which are respectively composed of right middle temporal gyrus and left middle temporal gyrus. They are correlated with a symmetrical pair rs-IC57 and rs-IC75, which are also replicated in a symmetrical pair rep-sIC25 and rep-sIC18. More interesting, structural components, s-IC40 and s-IC91, containing voxels in different sub-divisions of anterior cingulate cortex, are correlated with two components rs-IC39, rs-IC90 and replicated in rep-sIC 85, rep-sIC 15, centered on the same sub-divisions of anterior cingulate cortex.

### Cerebellar network

Component s-IC53 and s-IC46 respectively represent left and right cerebellum, which are correlated with rs-IC26 and rs-IC40, peaking at left and right cerebellum respectively, and replicated in nine components **(Figure S8D)**. Component s-IC36, primarily composed of vermis, is correlated with rsfMRI component rs-IC8 and replicated in rep-sIC1 and rep-sIC48, centered on vermis, a narrow midline zone in cerebellum. Structural components s-IC10 and s-IC11 are correlated with rs-IC26, replicated in rep-sIC53 and rep-sIC4, primarily comprised of bilateral cerebellum. Results from previous studies have found higher scores on vocabulary, reading, working memory and set-shifting were associated with increased GM in the posterior cerebellum (Moore DM et al. 2017), which is an example of how structural abnormalities directly relate to functional processing.

### Subcortical network

Subcortical structure s-IC3, comprising the putamen and parts of caudate, presents high S-F correspondence with functional component rs-IC33 in discovery dataset **(Figure S8E)**. The other subcortical component s-IC17, composed of bilateral caudate, is correlated with the same rsfMRI component, primarily comprising the bilateral putamen and caudate. The putamen and caudate contain the same types of neurons and circuits, which together form the dorsal striatum. The dorsal striatum plays an important role in the motor and reward systems, which receives inputs from cortical regions and then serves as the primary input to the rest of basal ganglia (Ferre S et al. 2010). Thus the putamen and caudate are likely to be decomposed in one functional component by ICA because of the similar brain function. However, as a white matter tract in the dorsal striatum structurally separates the caudate nucleus and the putamen (Ferre S *et al*. 2010), the putamen and caudate are more likely to be decomposed into two components by ICA for the structural data.

### Limbic network

Only component s-IC23, primarily composed of the hippocampus, has been revealed to be correlated to rs-IC44 and replicated in rep-sIC 77, primarily associated with memory function (van Strien NM et al. 2009). While S-F coherence of para-hippocampus was also observed in the discovery data as shown in **Figure S3-1**, the pair was not well replicated.

### Dorsal_attention network

Component s-IC67, which also largely covers the precuneus, is correlated to rs-IC30 and rs-IC35, replicated in five components **(Figure S9G)**. Two structural components s-IC 25 and s-IC27 are revealed to be correlated with the same functional component rs-IC76, which comprised of both parts of middle occipital gyrus and inferior parietal gyrus. Different from component s-IC79 and s-IC95, which also comprised of middle occipital but belongs the visual network, s-IC25 and s-IC27 are more close to the inferior parietal gyrus, sorted to the dorsal attention network. Inferior parietal gyrus is associated with bottom-up attention (Igelstrom KM and MSA Graziano 2017).

### Ventral_attention network

Structural component s-IC 30, centered at the supramarginal gyrus, is correlated with rs-IC 29 and replicated in rep-sIC73, which represents aspects of the supramarginal gyrus **(Figure S9H)**. Several studies have reported the supramarginal gyrus in interoceptive attention/awareness tasks (Kashkouli Nejad K et al. 2015). The other symmetrical components s-IC56 and s-IC76, are correlated with rs-IC31 and rs-IC94 respectively and replicated in rep-sIC24 and rep-sIC68, primarily composed of the insula.

### Frontoparietal network

The frontoparietal network yielded the lowest degree of S-F correspondence and the effects observed in the discovery sample were largely unreplicated **(Figure 3)**. Only component s-IC18, peaking in the superior frontal gyrus, is correlated to rs-IC 89 and rs-IC 85, and weakly replicates in rep-sIC 21, which represents activations over similar regions **(Figure S9I)**.

### Unimodel cortex exhibited better S-F correspondence than hetermodel cortex except default mode network

**Figure 5A** depicts the values of PC and MI of each replicated S-F pair in both datasets. Results show that components from somatomotor and visual networks present the highest S-F correspondence when adding up values from both metrics (PC and MI) and both cohorts, followed by default mode network (DMN), limbic and cerebellar networks. In contrast, the values in ventral_attention, dorsal_attention and frontoparietal networks are relatively lower than other networks, indicating more divergence between brain structure and function. Furthermore, we computed the S-F correspondence using PC or MI separately. As shown in **Figure 5B** and **Figure 5C**, either using PC or MI, the correspondences consistently show that the somatomotor, visual, and default mode network yield higher S-F correspondences, while components in the dorsal_attention and frontoparietal networks show more divergence between structure and function.

**Figure 5.**
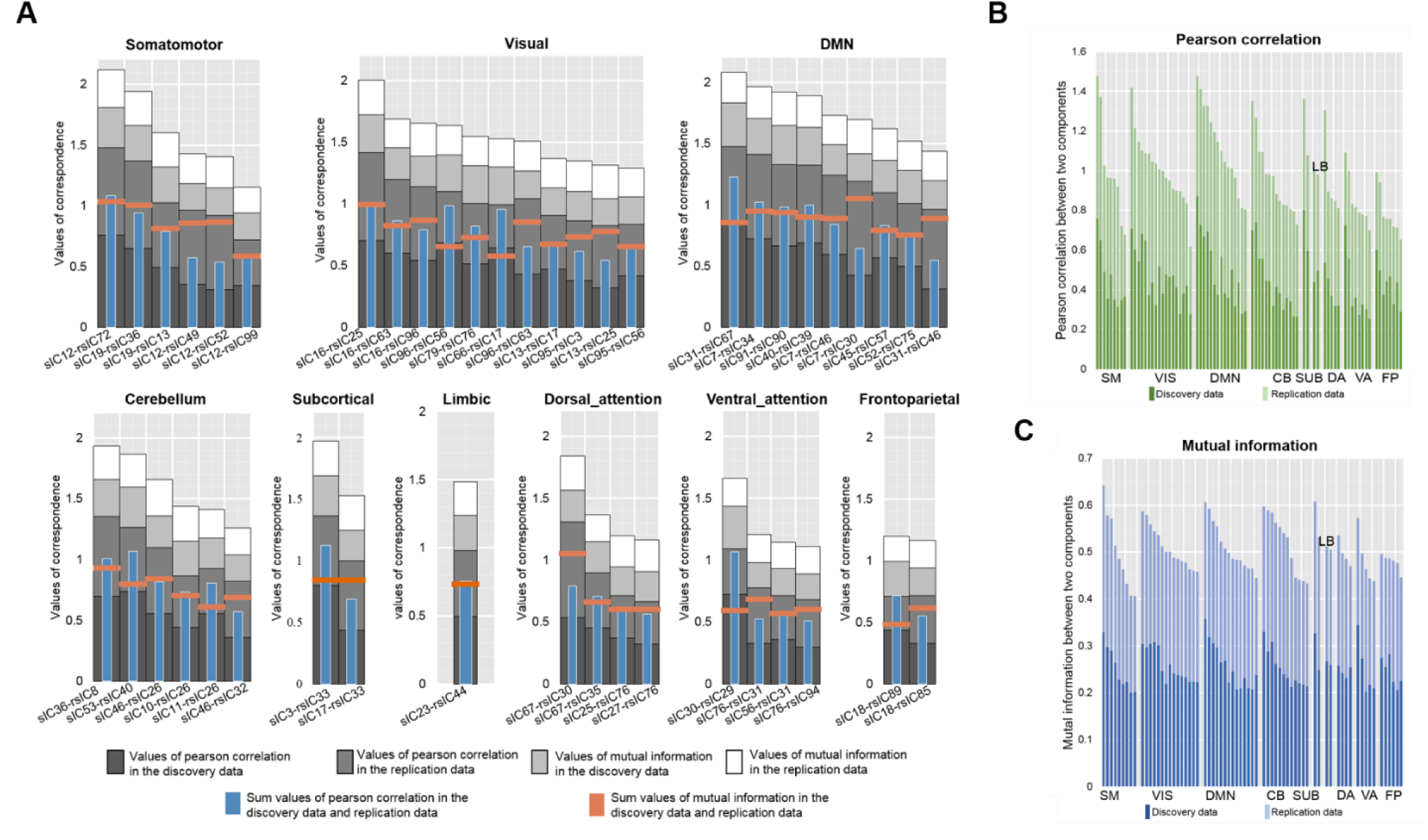
Comparison of S-F coherence across 9 domains. (A) The sum of Pearson correlation and mutual information in both discovery dataset and replication dataset of each component. The darker grey to white bars respectively represent the Pearson correlation in the discovery dataset, the Pearson correlation in the replication dataset, the mutual information in the discovery dataset and the mutual information in the replication dataset. The blue bar indicates the sum of Pearson correlation and mutual information from the discovery dataset, while the orange bar shows the sum of Pearson correlation and mutual information from the replication dataset. (B) The S-F correspondence in discovery and replication dataset measured only by Pearson correlation. The dark green bar represents the Pearson correlation in the discovery data and the light green color is the Pearson correlation in the replication data. (C) The S-F correspondence in discovery and replication dataset measured only by mutual information. The dark blue bar represents the mutual information in the discovery data and the light blue bar is the mutual information in the replication data.

## Discussion

In this study, we demonstrate that human structural architectures match intrinsic functional networks across the entire brain. With an ICA decomposition on gray matter volumes and spontaneous fluctuations of discovery data (6000+ scans), the structural components found are largely correspond to functional networks (62% matched 77.05%). Thus what we are seeing is the functional covariations in resting state BOLD signal, at the level of group networks across thousands of individuals, correlates in some cases with the structural correlations. To aid the robustness of the identified pairs, we replicated the S-F coherence in another independent dataset (UK Biobank 9000+ subjects). While 64.28% pairs are validated, the replicated percentage in each domain network are not identical. For example, replicated percentage in the frontoparietal network was 28.57%, implying that individual difference was large in this area. This was consistent with previous studies, which states that the frontoparietal network has emerged as the most distinctive fingerprinting feature for identifying individuals (Finn ES et al. 2015). In total, 45 S-F component pairs with high spatial consistency in both discovery and replication cohorts were identified, which were divided into 9 major networks, providing a stable S-F correspondence template that may be of use to the larger neuroimaing community.

The replicated S-F pairs and their correspondence values suggested that the unimodal cortical areas (somatomotor and visual network) show higher S-F correspondence, while heteromodal association cortical areas, especially the frontoparietal network, exhibit more divergence between intrinsic functional and structural networks. Unimodal cerebral cortex (Mendoza JE 2011) (somatomotor network and visual network), especially the precentral gyrus, supplementary motor area, and primary visual cortex (calcarine area, V1), exhibited higher S-F correpondence than other areas in both the discovery and replication dataset. The precentral gyrus and supplementary motor area are primarily associated with motor function (Graziano MS et al. 2002; Nachev P et al. 2008). Visual area V1 receives sensory inputs from the thalamus and plays an important role in the extraction of early visual features (Mechelli A et al. 2000). The somatomotor and visual regions have also been revealed to present the least individual differences in functional variability (Mueller S et al. 2013). However, the heteromodal association cortex (Green R 2004), except the DMN, show relatively low S-F correspondence (dorsal_attetntion network, ventral_attention network and frontoparietal network). The frontoparietal network presents the least S-F correspondence compared to the other networks. It has been previously demonstrated that the regions presenting the most prominent inter-subject variability are also the regions showing the most rapid expansion during human brain evolution (Mueller S *et al*. 2013), as well as greatest postnatal enlargement and a low maturation rate (Hill J et al. 2010). Wang *et al*. suggested that a greatly expanded and slowly maturing association cortex can provide a higher degree of freedom, both in physical space and time, for influences of environmental factors, potentially giving rise to inter-individual variability (Wang D and H Liu 2014), leading to weak S-F correspondences in these areas. In contrast, the unimodal regions mature early in life with a low expansion rate and are more stable to the environmental factors, which would present less inter-subject variability and more S-F correspondence.

The spatial consistence in default mode network is interesting; for example, the precuneus is split into two structural sub-regions (s-IC7 and s-IC67), which respectively correspond to two functional sub-regions (rs-IC34 and rs-IC30). The two sub-regions are consistent with previous subdivisions of the precuneus, which represent different functions : component s-IC7 and rs-IC34 are the posterior precuneus, which shows strong functional connectivity with visual-related areas, such as the cuneus (Margulies DS et al. 2009); component s-IC67 and rs-IC30 are the anterior precuneus, which exhibits strong functional connectivity with sensorimotor-related regions. Meanwhile, the other similar case is anterior cingulate cortex, where components s-IC40 and s-IC91 are correlated with two different anterior cingulate cortex in functional data (rs-IC39 and rs-IC90), consisting with previous subdivision of the anterior cingulate cortex (Bush G et al. 2000). Component s-IC40 is the dorsal part of the anterior cingulate cortex, which is connected with the prefrontal cortex and parietal cortex, participating in cognitive control (Shenhav A et al. 2016). By contrast, component s-IC91 is the ventral part of the anterior cingulate cortex, involving in generating emotional responses (Etkin A et al. 2011). Besides, structural components, s-IC45 and s-IC52, are symmetrical components, which are respectively composed of right middle temporal gyrus and left middle temporal gyrus, correlated with symmetrical pair rs-IC57 and rs-IC75 and replicated in a symmetrical pair rep-sIC25 and rep-sIC18. As shown in the Brainnetome Atlas (https://atlas.brainnetome.org/bnatlas.html), which yields functional characterization of sub-regions based on the BrainMap database using forward and reverse inferences (Fan LZ et al. 2016), the left middle temporal gyrus is primarily related to cognition, language and syntax, while the right middle temporal gyrus is involved in action and observation.

Despite the strengths of the current study, it still has some limitations. The first limitation is of course methodological: the structural images were generally collected at a much finer resolution (1 mm isotropic, generally) than the functional images (> 3 mm on a side), and the functional imaging is susceptible to signal drop-out in inferior areas or near the tissue border. We attempted to address this by reslicing the structural images to match the functional, and restricting the analysis to areas of gray matter, where signal in both modalities was robust. The second limitation is the lack of careful assessment of health status for the individuals included in discovery dataset. However, our results were further replicated in a large independent dataset with only healthy subjects. We believe the results may be more driven by the common characteristics on structural-functional correspondence. The third issue is that the MRI scanners and imaging protocols were not identical across the discovery and replication cohorts. However, since one important aim of this study is to identify a stable S-F template (i.e. a set of components with higher S-F correspondence), the replicated results in data with different scanners and imaging protocols provides a more generalizable result.

## Author contributions

V.D.C., J.S. and N.L. designed the experiments. N.L. conducted the analysis. V.D.C. and J.S. supervised and guided all aspects of the work. N.L., V.D.C. and J.S. wrote the paper. A.A. helped provide the information of the discovery data. Z.F. and E.D. helped select the components. D.C.G and A.L.R helped provide the information of the UK Biobank data. J.A.T, L.F., J.C., D.L., C.Z., D.C.G, A.L.R, M.T.B. and G.D.P. helped revise the paper. Describe the contributions of each author (use initials) to the paper.

## Data and materials availability

The multimodal data used in the present study can be accessed upon request to the corresponding authors.

## References

Abrol A, Damaraju E, Miller RL, Stephen JM, Claus ED, Mayer AR, Calhoun VD. 2017. Replicability of time-varying connectivity patterns in large resting state fMRI samples. Neuroimage. 163:160–176.

Alexander-Bloch A, Giedd JN, Bullmore E. 2013. Imaging structural co-variance between human brain regions. Nature Reviews Neuroscience. 14:322.

Alfaro-Almagro F, Jenkinson M, Bangerter NK, Andersson JLR, Griffanti L, Douaud G, Sotiropoulos SN, Jbabdi S, Hernandez-Fernandez M, Vallee E, Vidaurre D, Webster M, McCarthy P, Rorden C, Daducci A, Alexander DC, Zhang H, Dragonu I, Matthews PM, Miller KL, Smith SM. 2018. Image processing and Quality Control for the first 10,000 brain imaging datasets from UK Biobank. Neuroimage. 166:400–424.

Allen EA, Erhardt EB, Damaraju E, Gruner W, Segall JM, Silva RF, Havlicek M, Rachakonda S, Fries J, Kalyanam R, Michael AM, Caprihan A, Turner JA, Eichele T, Adelsheim S, Bryan AD, Bustillo J, Clark VP, Feldstein Ewing SW, Filbey F, Ford CC, Hutchison K, Jung RE, Kiehl KA, Kodituwakku P, Komesu YM, Mayer AR, Pearlson GD, Phillips JP, Sadek JR, Stevens M, Teuscher U, Thoma RJ, Calhoun VD. 2011. A baseline for the multivariate comparison of resting-state networks. Frontiers in systems neuroscience. 5:2.

Bassett DS, Bullmore ET. 2009. Human brain networks in health and disease. Curr Opin Neurol. 22:340–347.

Bush G, Luu P, Posner MI. 2000. Cognitive and emotional influences in anterior cingulate cortex. Trends Cogn Sci. 4:215–222.

Calhoun VD, Adali T, Pearlson GD, Pekar JJ. 2001. A method for making group inferences from functional MRI data using independent component analysis. Hum Brain Mapp. 14:140–151.

Cota Navin Gupta JAT, Vince D. Calhoun. 2017. Source Based Morphometry: Data-Driven Multivariate Analysis of Structural Brain Imaging Data. Brain Morphometry.

Du YH, Pearlson GD, Liu JY, Sui J, Yu QB, He H, Castro E, Calhoun VD. 2015. A group ICA based framework for evaluating resting fMRI markers when disease categories are unclear: application to schizophrenia, bipolar, and schizoaffective disorders. Neuroimage. 122:272–280.

Etkin A, Egner T, Kalisch R. 2011. Emotional processing in anterior cingulate and medial prefrontal cortex. Trends Cogn Sci. 15:85–93.

Fan LZ, Li H, Zhuo JJ, Zhang Y, Wang JJ, Chen LF, Yang ZY, Chu CY, Xie SM, Laird AR, Fox PT, Eickhoff SB, Yu CS, Jiang TZ. 2016. The Human Brainnetome Atlas: A New Brain Atlas Based on Connectional Architecture. Cerebral Cortex. 26:3508–3526.

Ferre S, Lluis C, Justinova Z, Quiroz C, Orru M, Navarro G, Canela EI, Franco R, Goldberg SR. 2010. Adenosine-cannabinoid receptor interactions. Implications for striatal function. Brit J Pharmacol. 160:443–453.

Finn ES, Shen XL, Scheinost D, Rosenberg MD, Huang J, Chun MM, Papademetris X, Constable RT. 2015. Functional connectome fingerprinting: identifying individuals using patterns of brain connectivity. Nat Neurosci. 18:1664–1671.

Graziano MS, Taylor CS, Moore T. 2002. Complex movements evoked by microstimulation of precentral cortex. Neuron. 34:841–851.

Green R. 2004. Heteromodal association cortex in schizophrenia. Am J Psychiatry. 161:1723–1724; author reply 1724.

Hill J, Inder T, Neil J, Dierker D, Harwell J, Van Essen D. 2010. Similar patterns of cortical expansion during human development and evolution. Proc Natl Acad Sci U S A. 107:13135–13140.

Igelstrom KM, Graziano MSA. 2017. The inferior parietal lobule and temporoparietal junction: A network perspective. Neuropsychologia. 105:70–83.

Kashkouli Nejad K, Sugiura M, Nozawa T, Kotozaki Y, Furusawa Y, Nishino K, Nukiwa T, Kawashima R. 2015. Supramarginal activity in interoceptive attention tasks. Neurosci Lett. 589:42–46.

Luo N, Sui J, Chen J, Zhang F, Tian L, Lin D, Song M, Calhoun VD, Cui Y, Vergara VM, Zheng F, Liu J, Yang Z, Zuo N, Fan L, Xu K, Liu S, Li J, Xu Y, Liu S, Lv L, Chen J, Chen Y, Guo H, Li P, Lu L, Wan P, Wang H, Wang H, Yan H, Yan J, Yang Y, Zhang H, Zhang D, Jiang T. 2018. A Schizophrenia-Related Genetic-Brain-Cognition Pathway Revealed in a Large Chinese Population. Ebiomedicine. 37:471–482.

Margulies DS, Vincent JL, Kelly C, Lohmann G, Uddin LQ, Biswal BB, Villringer A, Castellanos FX, Milham MP, Petrides M. 2009. Precuneus shares intrinsic functional architecture in humans and monkeys. Proc Natl Acad Sci U S A. 106:20069–20074.

Mechelli A, Humphreys GW, Mayall K, Olson A, Price CJ. 2000. Differential effects of word length and visual contrast in the fusiform and lingual gyri during reading. Proc Biol Sci. 267:1909–1913.

Mendoza JE. 2011. Unimodal Cortex. In: Kreutzer JS, DeLuca J, Caplan B, editors. Encyclopedia of Clinical Neuropsychology New York, NY: Springer New York p 2578–2578.

Misic B, Betzel RF, de Reus MA, van den Heuvel MP, Berman MG, McIntosh AR, Sporns O. 2016. Network-Level Structure-Function Relationships in Human Neocortex. Cereb Cortex. 26:3285–3296.

Moore DM, D’Mello AM, McGrath LM, Stoodley CJ. 2017. The developmental relationship between specific cognitive domains and grey matter in the cerebellum. Developmental cognitive neuroscience. 24:1–11.

Mueller S, Wang D, Fox MD, Yeo BT, Sepulcre J, Sabuncu MR, Shafee R, Lu J, Liu H. 2013. Individual variability in functional connectivity architecture of the human brain. Neuron. 77:586–595.

Nachev P, Kennard C, Husain M. 2008. Functional role of the supplementary and pre-supplementary motor areas. Nat Rev Neurosci. 9:856–869.

Power JD, Fair DA, Schlaggar BL, Petersen SE. 2010. The Development of Human Functional Brain Networks. Neuron. 67:735–748.

Segall JM, Allen EA, Jung RE, Erhardt EB, Arja SK, Kiehl K, Calhoun VD. 2012. Correspondence between structure and function in the human brain at rest. Frontiers in Neuroinformatics. 6.

Shenhav A, Cohen JD, Botvinick MM. 2016. Dorsal anterior cingulate cortex and the value of control. Nat Neurosci. 19:1286–1291.

Silver M, Montana G, Nichols TE, Neuroimaging AD. 2011. False positives in neuroimaging genetics using voxel-based morphometry data. Neuroimage. 54:992–1000.

Stephen Smith ED, Adrian Groves, Thomas E. Nichols, Saad Jbabdi, Lars T. Westlye, Christian K. Tamnes, Andreas Engvig, Kristine B. Walhovd, Anders M. Fjell, Heidi Johansen-Berg and Gwenaëlle Douaud. 2019. Structural variability in the human brain reflects fine -grained functional architecture at the population level. The Journal of Neuroscience.

Sui J, Qi S, van Erp TGM, Bustillo J, Jiang R, Lin D, Turner JA, Damaraju E, Mayer AR, Cui Y, Fu Z, Du Y, Chen J, Potkin SG, Preda A, Mathalon DH, Ford JM, Voyvodic J, Mueller BA, Belger A, McEwen SC, O’Leary DS, McMahon A, Jiang T, Calhoun VD. 2018. Multimodal neuromarkers in schizophrenia via cognition-guided MRI fusion. Nat Commun. 9:3028.

van Strien NM, Cappaert NLM, Witter MP. 2009. The anatomy of memory: an interactive overview of the parahippocampal-hippocampal network. Nature Reviews Neuroscience. 10:272–282.

Wang D, Liu H. 2014. Functional Connectivity Architecture of the Human Brain: Not All the Same. The Neuroscientist. 20:432–438.

Xu L, Groth KM, Pearlson G, Schretlen DJ, Calhoun VD. 2009. Source-Based Morphometry: The Use of Independent Component Analysis to Identify Gray Matter Differences With Application to Schizophrenia. Hum Brain Mapp. 30:711–724.

Yeo BTT, Krienen FM, Sepulcre J, Sabuncu MR, Lashkari D, Hollinshead M, Roffman JL, Smoller JW, Zoller L, Polimeni JR, Fischl B, Liu HS, Buckner RL. 2011. The organization of the human cerebral cortex estimated by intrinsic functional connectivity. J Neurophysiol. 106:1125–1165.

